# Residual-matching: An efficient alternative to random sampling in human subjects research recruitment

**DOI:** 10.1101/2020.04.14.041384

**Authors:** Andrew Hooyman, Matthew J. Huentelman, Sydney Y. Schaefer

**Affiliations:** Department of Biological and Health Systems Engineering, Arizona State University, Tempe, Arizona, USA; The Translational Genomics Research Institute, Neurogenomics Division, Phoenix, Arizona, USA

**Keywords:** Citizen Science, Recruitment, Simulation, Resource efficiency

## Abstract

Given the time- and resource-intense nature of human subjects research, we have developed a more intelligent approach to participant recruitment above and beyond random sampling that leverages pilot or preliminary results to reduce the overall number of participants needed for recruitment from an existing electronic cohort or database. Using open-access data from the General Social Survey (GSS) of the National Opinion Research Center, we generated pilot and validation datasets through a simulation to establish moderate and weak relationships based on linear regression. We then compared the performance of our residual-matching method against random sampling in their probabilities of achieving a given level of statistical power as well as their prediction accuracies. Results showed that the residual-matching method was superior to random sampling, yielding smaller sample sizes with equivalent mean square error. We therefore advocate the use of residual matching when scaling up pilot studies to conserve time and resources in larger follow-up studies.

## INTRODUCTION

In practically every human subjects research study, both physical resources (e.g., equipment, space) and temporal resources (e.g., scientists’ time, participants’ time) are necessary. Moreover, these resources are undoubtedly finite and limited, yet the expectation within the scientific community is to achieve results that are well-powered and generalizable, i.e. experiments with sufficient sample sizes for the given research question^1^. This therefore often creates a race to achieve sufficient statistical power before running out of time, money, equipment, or eligible participants. As a result there has been a renewed focus on how to best utilize resources, prevent waste, yet increase rigor in research^2,3^. This is especially true where large amounts of data are needed to reproduce or extend previous findings^4^. Imagine having just completed a small pilot study that showed a significant relationship between independent variable *A* (such as a history of smoking) and dependent variable *B* (like memory), but the study itself was underpowered and from a fairly homogenous (white non-Hispanic) sample. How now would one recruit to more rigorously test this hypothesis in a larger and more diverse sample, perhaps in Latinx participants in a way that is resource- and time-efficient?

Traditionally, participant recruitment for human subjects research would be based off of random sampling with some inclusion/exclusion criteria (e.g., adults over age 65 years, a history of smoking, etc)^5^. Then, once consented and enrolled, the study would commence, either by sending test materials (e.g., survey) to them to complete at home or by bringing the participant into a laboratory space to measure the relationship between a history of smoking and memory within each individual at random. Recently, studies have utilized citizen science (defined as at-large “public participation in scientific research”)^6^ and crowd-sourcing to recruit large numbers of participants at random^7,8^. However, studies as these that utilize random recruitment can be especially problematic since the resultant sample distribution is often non-random, meaning the study sample does not represent that of the intended population^9^. Our premise is that random sampling may prove to be inefficient and arguably wasteful, as some of the test materials or data collection slots will undoubtedly be given to outliers, or lead to an oversampling of a particular demographic that is not as sensitive to the hypothesized relationship.

We propose a solution to utilize strategic recruitment methods when using a database of eligible participants, such as a large electronic cohort who have agreed to be contacted for future research studies and report a history of smoking. This statistical approach identifies individuals who would be most sensitive to the underlying relationship being investigated, based on other characteristics. This approach also aids in refining one’s hypothesis. Continuing with the example above, if a set of researchers aims to test whether their previous finding of a relationship between smoking history and memory in one racial group generalizes to another racial group, they would want to match their second study’s recruitment with the exact age, sex and education distribution of the participants in the initial study. However, it is rare that factors like age, sex, and education have equal and uniform effects on the relationship of interest, so simply age-, sex-, and education-matching one’s groups may not be sufficient. Furthermore, there may also be unexpected participant characteristics, such as a family history of dementia, which could be important to evaluate. We have therefore developed a method that can be used to first identify the participants in a pilot sample who were most sensitive to the relationship being tested (e.g., smoking history vs. memory), and then generate a list of additional participant characteristics of future study participants based on those who were most sensitive. These then would be the individuals within the database that the research team would first want to recruit. We note, however, that this approach should *not* be seen as ‘cherry-picking’, where one may try to identify individuals who are biased towards a preferred or favorable hypothesis. Instead, our method (as detailed below) is data-driven and utilizes matching methods that remove experimenter bias^10^.

The aim of this paper is to outline and test a simulation to compare how our distribution matching method termed *residual-matching* replicates an established finding with a specific level of power relative to random sampling for participant recruitment. Results of the simulation will provide context of what percentage of resources are saved or lost between each method. Ideally, our method will demonstrate a lower number of participants needed to achieve a certain level of power without a loss in prediction accuracy.

## METHODS

### Dataset used in simulation

We applied our recruitment simulation on an open-source dataset found here https://vincentarelbundock.github.io/Rdatasets/datasets.html. Specifically, we are using the *GSSvocab* dataset that is from the General Social Survey (GSS) of the National Opinion Research Center of the University of Chicago. This dataset has 28,867 observations on the follow eight variables: year, gender, native born, age in years, age by decade, education years, education by group (<12, 12, 13-15, 16, and >16 years of education) and vocabulary, measured as the number of words out of 10 correct on a vocabulary test.

### Generating pilot and validation datasets and establishing relationships between variables

With the GSSvocab dataset, we generated a simulation with an outer and inner loop. The outer loop of the simulation randomly divided the entire GSSvocab dataset into pilot and validation/electronic cohort datasets. Then, an initial pilot relationship was performed using linear regression to predict the number of correct vocabulary words based on sex, age, and years of education. Next, the inner loop iteratively performed a replication of the pilot relationship across a range of participant sampling (each iteration of the inner loop performs the replication with a larger and larger participant number), with both recruitment methods on the validation dataset. Then, achieved power and prediction error of the replicated regression model on the pilot data were stored for each iteration as simulation outcome variables. Once the inner loop was complete, the outer loop repeated 10 times, initiating a new variation of the pilot relationship that the inner loop then tested across a range of sample sizes dependent on the strength of relationship analyzed (see below). We performed the simulation twice with both a known moderate and known weak relationship to vocabulary (shown here) to demonstrate the robustness of the “residual-matched” (RM) method:

- Moderate relationship: vocab ~ age + sex + education; R2 =. 24; tested over range of sample sizes: 20 – 100
- Weak relationship: vocab ~ age + sex; R^2^ =. 003; tested over range of sample sizes: 1,000 - 10,000

### Recruitment methods: Residual-matched vs. random sampling

#### Identification of pilot participants most sensitive to the established relationship

Our proposed method requires a known existing relationship, either from previous studies (i.e., publicly available data from open-access databases) or pilot work, to determine covariate distributions that will then be applied to our strategic recruitment method, which we call the ‘residual-matched’ or RM method. In the first simulation using the GSSvocab dataset, we used linear regression to predict the number of correct vocabuary words based on age in years, education in years and sex (which yielded a moderate relationship), and in the second simulation, we predicted the number of correct vocabulary words by age and sex only (which yielded a weak relationship). All analyses are performed in R version 3.5.5^11^.

From the resulting regression model of this initial pilot set (the size, N, of which is determined by the maximum level of matching we wished to do in the simulation), we sorted the data based on each pilot participant’s linear regression residual, with the lowest residual sorted to the top (indicating individuals most sensitive to the given relationship), and the highest residual sorted to the bottom (indicating individuals who were furthest from the established relationship). Then given *a select amount of resources*, *X* number of individuals from the residual-sorted dataset were identified. These individuals were then matched to participants in the validation set based on by sex, age in years and education in years, using the MatchIt algorithm^12^ (described next).

#### Application of each recruitment method to a validation dataset

The MatchIt algorithm is the implementation of a nonparametric data preprocessing method that assigns a dichotomous grouping between two sets of data with the same dependent variables. In our use of the MatchIt algorithm, we assigned one variable to our pilot dataset and then had MatchIt identify the nearest match of each pilot participant in the electronic cohort based on age, sex and years of education. The matched samples from the electronic cohort were then assigned their own grouping variable, which effectively created the list of validation participants to be recruited in the experiment for replication of the previous pilot relationship. In parallel, as an implementation of the random sampling method, *X* number of participants were randomly sampled from the same electronic cohort as the RM method.

#### Simulation outcome variables: Replication results of each recruitment method

Linear regression was used to predict the number of correct vocabulary words using age, sex and years of education with both the RM and random sampling validation datasets. The power achieved by each regression analysis was calculated and stored. Additionally, the resulting regression models were also used to predict the previous pilot dataset and mean squared error between the predicted values of the regression analysis and actual values of the pilot dataset calculated, and were also stored for further analysis.

### Statistical analysis of simulation results

Logistic regression was utilized to determine effectiveness of each method to achieve a specific level of power given a certain sample size. Specifically, logistic regression was utilized to determine the probability of achieving a power level of. 8 between each method based on varying sample size. An independent t-test was utilized to determine mean differences between the recorded mean squared error of each recruitment method.

## RESULTS

### Probability of achieving specific power level given sample size

#### For the moderate relationship (vocab ~ age + sex + education; R^2^ =. 24)

Results of our logistic regression demonstrated a significant difference in the probability of each method (RM vs random sampling) achieving a power of. 8 or greater given a specific sample size (p <. 05). Results demonstrated that the RM method achieved a greater than 50% probability of achieving a power of. 8 with a sample size of 44 compared to the random method that required a sample size of 54 (Fig 1).

**Figure 1.**
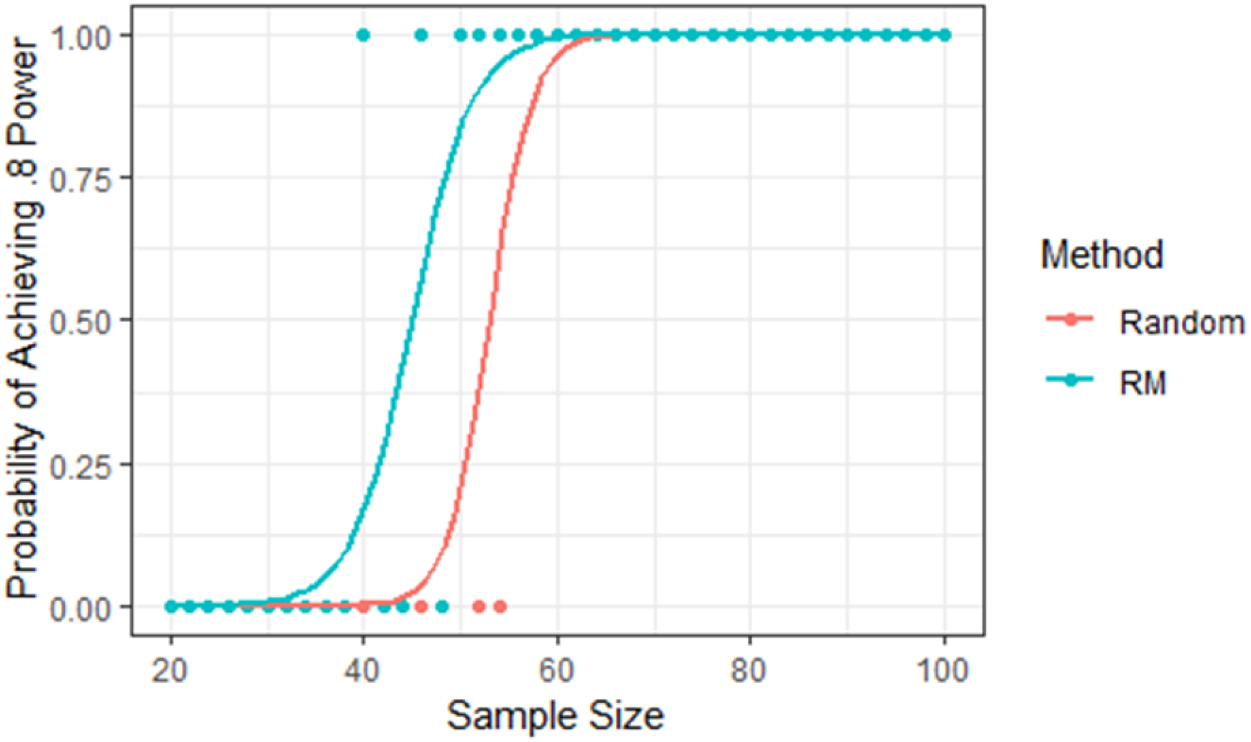
Moderate Relationship - Probability of Achieving Power of. 8 Based on Sample Size and Method. Based on a range of smaple sizes the RM method demonstrates a greater probability of achieving statistical power of. 8 with a smaller sample size than the Random sampling method.

#### For the weak relationship (vocab ~ age + sex; R^2^ =. 003)

Results of our logistic regression demonstrated a significant method difference in the probability of each method (RM vs random sampling) achieving a power of. 8 or greater given a specific sample size (p <. 05). Results demonstrated that the RM method achieved a greater than 50% probability of achieving a power of. 8 with a sample size of 3000 compared to the random method that required a sample size of 4500 (Fig 2). It is noted that the relative effectiveness of the RM method compared to the random sampling in both the moderate (54:44, or 1.2 times) and weak (4500:3000, or 1.5 times) relationships was similar.

**Figure 2.**
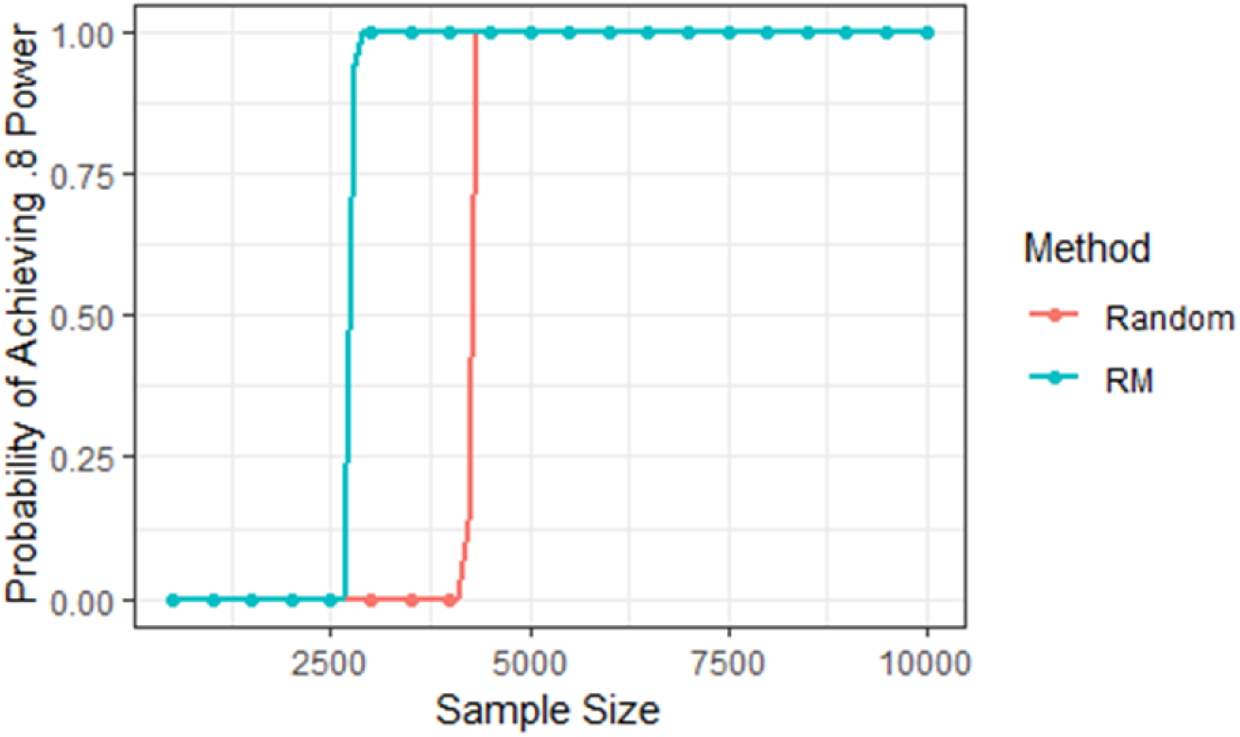
Weak Relationship - Probability of Achieving Power of. 8 Based on Sample Size and Method. Based on a range of smaple sizes the RM method demonstrates a greater probability of achieving statistical power of. 8 with a smaller sample size than the Random sampling method.

### Validation dataset prediction error

It is plausible that although the RM method performs better than random sampling in terms of probability of achieving a given level of power, it may perform worse in terms of prediction error. However, results from our independent t-test indicated that mean squared error (by which the cohort regression model predicted the original pilot data) was not significantly different between the RM and random sampling methods (p >. 05, Fig 3). This demonstrates that overall robustness of the RM method is equivalent to that of the random sampling method, achieving the same model accuracy with a lower sample size. We therefore consider this approach to be a more responsible and efficient method of recruitment as it can conserve a number of resources both at the start of a study and throughout.

**Figure 3.**
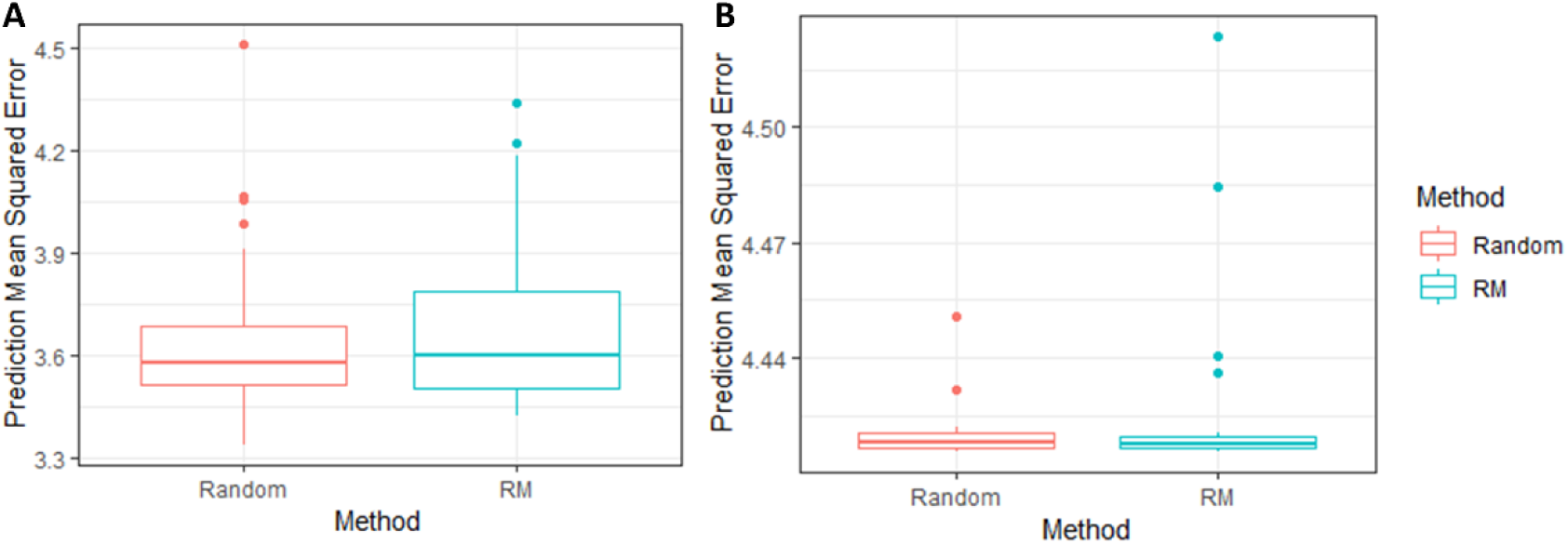
Mean Squared Error of Cohort Model Predictions on Original Pilot Data by Method A. Moderate Realtionship, B. Weak Relationship. Both recuitment methods achieved a similar mean squared error i.e. pridiction accuracy, on the original pilot data from their respective cohort regression models.

## DISCUSSION

Results from this experiment showed that our residual-matched (RM) method was capable of achieving a given level of statistical power with a smaller N compared to random sampling, without loss of prediction accuracy in both moderate and weak relationships. Overall, the RM method required approximately 30% fewer participants than random sampling. This provides a strong case for the application of this method in ‘citizen science’ or crowdsourcing designs that recruiting from electronic cohorts (i.e., MindCrowd,^13^) with the aim of replicating or generalizing an existing or pilot relationship. This also ‘closes the loop’ proposed recently by Huentelman et al. (2020) regarding how electronic cohorts and pilot testing can build off each other for hypothesis testing and generation^14^ in a more efficient way (Fig 4). The RM method may be attractive for performing well-powered research with a limited amount of resources (time, money, or equipment). Conserving resources within human subjects research also can accelerate the development and implementation of follow-up studies, since less data are needed to complete a given project. With more competition for funding and larger budget cuts, it is critical that researchers be cognizant of how to do more with less. Overall, we have demonstrated that our residual matching (RM) method, which is easy to implement, is capable of intelligent participant recruitment for human subjects research for the purposes of well-powered research with fewer participants than random sampling.

**Figure 4.**
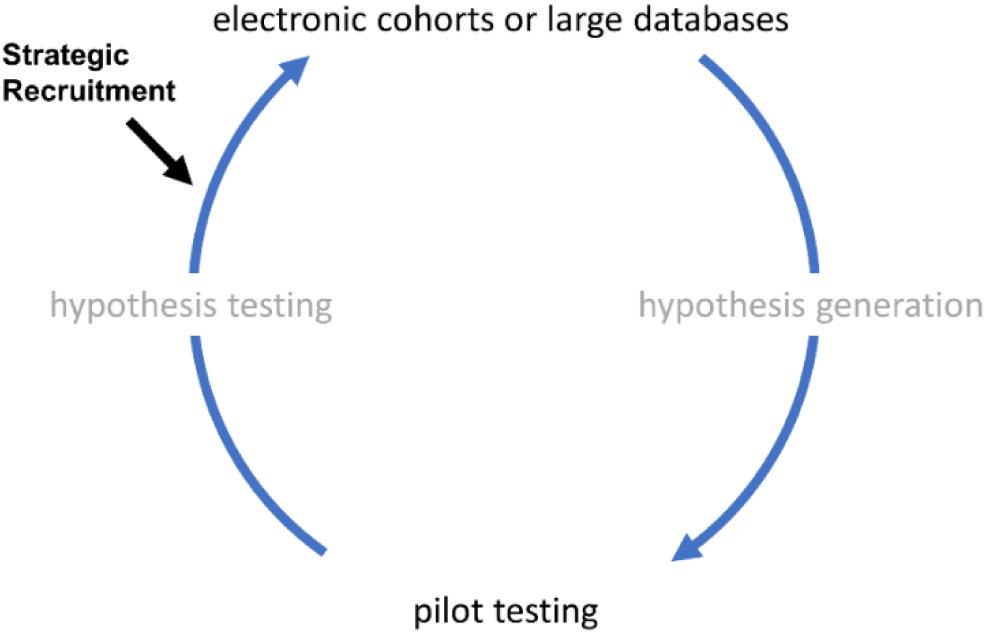
The Huentelman et al. Model for Reinventing Research in the Digital Age. Within this model pilot testing and electronic cohorts inform each other in a cyclical nature. We demonstrate how insertion of strategic recuitment may reduce waste and improve resource allocation in the conversion of pilot testing into large scale research.

### Limitations

The major limitation of the RM method is that it requires a known relationship of good quality (i.e., analysis assumptions met and no outliers present) to identify sensitive individuals through residual matching. If the pilot relationship is based on the presence of a few outliers (i.e., individuals who apply significant leverage to the mode), then the RM method will be biased toward those outliers. It is noted, however, that random sampling would not solve this issue either, and could in fact waste even more time and resources recruiting for an effect that is not really present. It is in these cases, in general, that diligent and ethical decisions need to be made on whether to even proceed to a larger scale study on these initial findings at all^15,16^. Another current limitation is the availability of a robust electronic cohort or dataset that meets the needs of one’s research question at hand. However, with the continued push for data sharing in both clinical trials and open source repositories, it has become easier to perform preliminary analyses that could utilize the RM method. We advocate for, whenever possible, testing one’s initial pilot findings with relevant independent datasets, such as those from any of the open-access datasets in the Inter-University Consortium for Political and Social Research. In addition, we promote the development and utility of such datasets to further generate new hypotheses that can be tested in small pilot studies (Fig. 4). In short, this paper proposes an easy, data-driven approach for recruiting human subjects for research in a time- and resource-efficient manner.

